# RiboTag-based RNA-Seq uncovers oligodendroglial lineage-specific inflammation in autoimmune encephalomyelitis

**DOI:** 10.1101/2024.12.24.630168

**Authors:** Yuhang Wang, Sudeep Ghimire, Ashutosh Mangalam, Zizhen Kang

## Abstract

Oligodendroglial lineage cells (OLCs) are critical for neuronal support functions, including myelination and remyelination. Emerging evidence reveals their active roles in neuroinflammation, particularly in conditions like Multiple Sclerosis (MS). This study explores the inflammatory translatome of OLCs during the early onset of experimental autoimmune encephalomyelitis (EAE), an established MS model. Using RiboTag-based RNA sequencing in genetically modified Olig2-Cre RiboTag mice, we identified 1,556 upregulated and 683 downregulated genes in EAE OLCs. Enrichment analysis indicated heightened immune-related pathways, such as cytokine signaling, interferon responses, and antigen presentation, while downregulated genes were linked to neuronal development and myelination. Notably, OLCs expressed cytokines/chemokines, and their receptor, highlighting their active involvement in neuroinflammatory signaling. Functional studies demonstrated that interferon-gamma (IFN-γ) signaling in OLCs exacerbates EAE pathology by enhancing antigen presentation and chemokine production, whereas interferon-beta (IFN-β) signaling showed minimal impact. These findings provide novel insights into the inflammatory role of OLCs in EAE and suggest therapeutic potential in targeting OLC-mediated neuroinflammation for MS and related disorders.

## Introduction

Oligodendrocyte Progenitor/Precursor Cells (OPCs) constitute the fourth largest population of glial cells within the central nervous system (CNS), accounting for approximately 5% of all CNS cells and enduring from infancy into adulthood ^1, 2^. Remarkably, OPCs exhibit an even distribution throughout the CNS, lacking specific niches and distinguishing themselves from transient cell populations typical of other progenitor cells. Traditionally recognized for their role in generating mature oligodendrocytes (mOLs) responsible for axon myelination in the spinal cord and brain, OPCs originate from radial glia in ventricular zones during embryonic development. Subsequently, they undergo local proliferation, migration, and differentiation into mOLs during both development and following injury ^3–5^. Given their persistent presence across the CNS, it has been hypothesized that OPCs may serve functions beyond mere precursor roles over extended periods. Indeed, emerging research has collectively revealed myelination-independent functions of OPCs in both health and disease contexts, including involvement in inflammatory responses, neural circuitry remodeling, axon regeneration, angiogenesis, and neuronal development ^6, 7^.

Multiple sclerosis (MS) is an autoimmune demyelinating disorder impacting 2-3 million individuals worldwide, with a prevalence ranging from 50 to 300 per 100,000 people ^8–10^. Although the exact mechanisms remain elusive, it is well-established that mOLs are targeted in MS pathology. The inflammatory milieu within the central nervous system is considered a primary driver of demyelination in MS and impedes remyelination by promoting mOLs death while hindering the maturation and proliferation of OPC ^2^. Intriguingly, recent findings suggest that OPCs not only respond to inflammatory signals but actively participate in modulating inflammation, influencing disease progression ^11–13^. Our research has demonstrated that NF-κB activator 1 (Act1)-mediated IL-17 signaling in OPCs plays a critical role in the pathogenesis of experimental autoimmune encephalomyelitis (EAE), an animal model for MS. IL-17 not only inhibits OPC maturation and promoting its apoptosis, but also induces the production of inflammatory cytokines and chemokines, thereby exacerbating neuroinflammation ^14^. Furthermore, OPCs aggregate in the perivascular regions of active MS lesions, contributing to blood-brain barrier disruption and CNS inflammation ^15^. Lipoprotein-related protein 1 (LRP1) in OPCs has been implicated in modulating neuroinflammation through antigen cross-presentation ^16^. Additionally, studies have identified expression of genes involved in antigen processing and presentation by oligodendroglia cells in human MS tissues ^17^ and MS mouse models ^18, 19^, including MHC-I and II genes. Notably, single-cell RNA sequencing of the oligodendroglia lineage during the peak of EAE in mice revealed a diverse immune profile present in both OPCs and Ols ^18^, highlighting the potential for varied immune functions within this lineage in the context of MS ^20^.

While single-cell transcriptomic analysis presents an opportunity to explore the full spectrum of oligodendroglial cells and their transcriptome phenotypes, this approach does come with certain limitations. Detecting abundant transcripts can be challenging due to the simultaneous sequencing of numerous cell types, and the process of isolating single cells through sorting may potentially alter gene expression ^21, 22^. Ribosome tagging (RiboTag) technology has emerged as a valuable in vivo method for studying gene expression and mRNA translation in specific cell types that are difficult and time-consuming to isolate using traditional techniques ^21, 23^. This method involves using a RiboTag mouse line with a modified Rpl22-HA (hemagglutinin) allele, which can be activated by Cre recombinase. Consequently, immunoprecipitation of HA enables the purification of polysomes and associated mRNAs from the target cells, ensuring that the isolated mRNA represents actively translated proteins since only ribosome-bound RNAs are purified ^24^. In order to analyze the specific transcriptome of the oligodendroglial lineage in the EAE model, RiboTag mice were crossed with Oligo2-Cre transgenic mice ^25^ to induce HA expression exclusively in oligodendroglial lineage cells (refer to as OLCs). Subsequently, mRNA from the oligodendroglial lineage in the spinal cords was sequenced and subjected to bioinformatic analysis to identify cell-specific gene expression patterns.

In this study, we identified 1,556 upregulated and 683 downregulated genes in OLCs from EAE mice. Enrichment analysis showed increased immune pathways, such as cytokine signaling, interferon responses, and antigen presentation, while downregulated genes were associated with myelination and neuron development. Notably, OLCs expressed cytokine, chemokine, and toll-like receptors. Further testing indicated that IFN-γ signaling in OPCs exacerbates EAE pathology, likely by enhancing antigen processing, presentation, and cytokine/chemokine production, whereas IFN-β signaling appeared non-essential for disease progression. These findings underscore a pathogenic role for IFN-γ signaling in OLCs during EAE-related inflammation.

## Materials and methods

### Mice

Ifngr1 floxed mice ^26^ (referred to as Ifngr1^fl^, JAX 025394), Ifnar1^fl^ mice ^27^(JAX 028256), Olig2-cre mice ^25^(JAX 025567), Pdgfra-cre^ER^ mic e^28^ (JAX 018280),Rpl22-HA flexed mice (refer to as RiboTag mice) ^24^, and C57BL/6J mice (JAX 000664) were all procured from Jackson Laboratory. To induce PDGFRa expression in Pdgfra-CreER mice, tamoxifen was administered at a dose of 1 mg/mouse/day via intraperitoneal (i.p) injection for 5 consecutive days. The experimental mice, aged 8-12 weeks, included both males and females and were housed in individually ventilated cages under specific pathogen-free conditions within accredited animal facilities. All animal procedures were ethically approved by the Institutional Animal Care and Use Committee of the University of Iowa.

### Induction and assessment of EAE

The mice were immunized by subcutaneous injection in the right and left flank with 200 μg of MOG35-55 peptide emulsified in a 1:1 ratio in complete Freund’s adjuvant (CFA) supplemented with 5 mg/ml Mycobacterium tuberculosis (Becton Dickinson, Franklin Lakes, NJ). Additionally, mice immunized with myelin peptides were each administered 500 ng of pertussis toxin (List Biologicals, Campbell, CA) via intraperitoneal injection on days 0 and 2. Clinical disease scores were documented daily based on the following criteria: 0 for no symptoms; 1 for tail weakness or loss of tail rigidity; 2 for mild hindlimb weakness; 3 for partial hindlimb paralysis; 4 for complete hindlimb paralysis; and 5 for moribund status or death, as previously described ^14, 29^.

### Histological analysis

The spinal cords were harvested from PBS-perfused EAE mice 25 days after EAE induction, approximately corresponding to the peak of disease severity. For paraffin sections, the spinal cords were fixed in 10% formalin, and all sections used were 10 μm thick. These paraffin sections were stained with hematoxylin and eosin (H&E) to assess inflammation and demyelination. For frozen sections, fresh spinal cords were embedded in OCT (Tissue-Tek) and rapidly frozen in liquid nitrogen. The frozen sections, also 10 μm thick, were utilized for fluorescent staining. These sections were stained with specific antibodies: anti-CD3 (Abcam, ab16669), anti-MBP (Cell Signaling Technology, CST#788965), Anti-GFAP (DSHB, University of Iowa, Cat# N206A/8), anti-HA (Sigma Aldrich, # H9658 or CST #3724), anti-NeuN (CST, #24307), or anti-GST-π (ADI-MSA-102-E, Enzo Life Science). The nuclei were counter-stained with DAPI. Antigen visualization was achieved through incubation with fluorescence-conjugated secondary antibodies from Molecular Probes.

### Quantitative real-time PCR (qRT-PCR)

Real-time PCR analysis was conducted to measure gene expression levels. RNA was extracted from spinal cord tissue or cultured OPCs using TRIzol (Invitrogen) as per the manufacturer’s instructions. The expression levels of all genes are presented as arbitrary units relative to the expression of the β-actin gene. Subsequently, the extracted RNA was promptly reversed transcribed into complementary DNA (cDNA). Quantitative PCR (Q-PCR) analysis was carried out using SYBR Green Real-time PCR Master Mix on a Real-Time PCR System (Applied Biosystems).

### Flow cytometry analysis

Oligodendrocyte precursor cells (OPCs) were cultured from Ifngr1^fl/fl^mice, olig2-cre Ifngr1^fl/fl^mice, and Pdgfra-cre^ER^ Ifnar1^fl/fl^ mice. OPCs from Pdgfra-cre^ER^ Ifnar1^fl/fl^ mice were treated with either 1μM 4-hydroxytamoxifen in the last three days of culture to induce Cre expression or corn oil vehicle as a control. Subsequently, OPCs were stained with fluorescence-conjugated antibodies targeting CD140a (Biolegend, Clone APA5), in combination with either anti-IFNGR (Biolegend, Clone 2E2) or anti-IFN-AR (Biolegend, Clone MAR1-5A3) antibodies, all diluted at a 1:100 ratio when used. The stained cells were analyzed using a Cytek Aurora cytometer, and the data were further analyzed with FlowJo software.

### Primary culture of oligodendrocyte progenitor cells (OPCs)

The culture of oligodendrocyte precursor cells (OPCs) was conducted following previously established protocols ^14, 30^. Briefly, neurospheres were prepared from E14.5 embryos obtained from timed pregnant females. The neurosphere medium consisted of DMEM/F12, B27 neuronal supplement (Invitrogen), and 10 ng/ml recombinant mouse epidermal growth factor (EGF, R&D Systems). Floating neurospheres were passaged at a 1:3 ratio in the same medium every 3–4 days. To obtain pure OPCs, neurospheres were dissociated after 2–3 passages, and the dissociated cells were plated on poly-D-lysine-coated plates under the same neurosphere medium, but with 10 ng/mL fibroblast growth factor (FGF) and 10 ng/mL platelet-derived growth factor (PDGF) (Peprotech) instead of EGF. Following 6 days of culture, approximately 90% of cells were CD140a-positive (CD140a+), confirming their identity as OPCs. The cultured OPCs were utilized for genotyping or functional studies as detailed in the figure legend.

### RiboTag immunoprecipitation

RiboTag immunoprecipitations were performed as described previously ^24^, with some modifications. Initially, the spinal cords from EAE mice at the onset of disease and naïve control mice were homogenized using a tissue homogenizer in a supplemented homogenization buffer composed of 5% (w/v) concentration, containing 1% NP-40, 100 mM KCl, 100 mM Tris-HCl pH 7.4, and 12 mM MgCl_2_. This buffer was further enriched with 100 μg/ml cycloheximide, a protease inhibitor cocktail, 1 mg/ml heparin, RNase inhibitors (Promega, Cat#N2115), and 1 mM dithiothreitol (DTT). The resulting homogenates underwent centrifugation, and the supernatant was then incubated at 4°C for 4 hours with 5 μl of anti-HA antibody to capture the HA tagged ribosomes (CST, Rb anti-HA #3724, 1:200). Subsequently, Pierce Protein A/G Magnetic Beads (Thermo Fisher, Cat#88803) were bound to the antibody-ribosome complex through overnight incubation at 4°C on a rotator. The beads were then subjected to three washes with a high salt buffer comprising 300 mM KCl, 1% NP-40, 50 mM Tris-HCl pH 7.4, 12 mM MgCl2, 100 μg/ml cycloheximide, and 500 μM DTT. RNA was released from the ribosomes using 350 μl of RLT buffer (from Qiagen RNeasy kit) with 1% 2-mercaptoethanol. Subsequent RNA purification was performed using the RNeasy Plus Micro kit (Qiagen 74034) following the manufacturer’s instructions, and the eluted RNA was collected in 16 μl of RNase-free water and stored at −80°C. Additionally, for each sample, 50 μl of homogenate (prior to the addition of anti-HA antibody) was set aside, stored at −20°C overnight, and purified using the RNeasy Micro kit as an ‘input’ sample to determine oligodendroglial RNA enrichment. The RNA integrity number (RIN) was determined using the Agilent RNA 6000 Pico Kit (Agilent Technologies, Cat#5067-1513).

### RNA-Seq

We initiated RNA sample preparations using 1 μg of RNA per sample. Libraries were crafted employing the NEBNext Ultra RNA Library Prep Kit for Illumina (NEB), following the manufacturer’s guidelines, with index codes appended to identify sequences for each sample. Initially, mRNA was isolated from total RNA using poly(T) oligo-attached magnetic beads. Fragmentation ensued using divalent cations at elevated temperatures in NEBNext First Strand Synthesis Reaction Buffer (5x). First-strand cDNA synthesis employed a random hexamer primer and M-MuLV reverse transcriptase (RNase H). Subsequent second-strand cDNA synthesis utilized DNA polymerase I and RNase H. Remaining overhangs were converted to blunt ends via exonuclease–polymerase activities. Following adenylation of the 3′ ends of DNA fragments, NEBNext Adaptor with a hairpin loop structure was ligated to facilitate hybridization. To preferentially select cDNA fragments around 150–200 bp in length, library fragments were purified using the AMPure XP system (Beckman Coulter). Subsequently, 3 μl USER Enzyme (NEB) was applied to size-selected, adaptor-ligated cDNA at 37 °C for 15 minutes followed by 5 minutes at 95 °C before PCR. PCR employed Phusion High-Fidelity DNA polymerase, Universal PCR primers, and Index (X) Primer. Finally, PCR products were purified using the AMPure XP system, and library quality was assessed on the Agilent Bioanalyzer 2100 system. Index-coded sample clustering was executed on the Illumina NovaSeq sequencer per the manufacturer’s instructions. Following cluster generation, libraries were sequenced on the same machine, yielding paired end reads.

### RNA-Seq data analysis

Paired reads underwent initial processing on the University of Iowa high-performance computing cluster for sequence quality control and analysis. Adapters were initially removed and bases with a Phred score <20 were trimmed from both forward and reverse reads by using Metawrap read_qc module from Metawrap package (version 1.3.2) ^31^. The quality-controlled fastq files were then aligned to Mus_musculus.GRCm39.cdna.all.fa (downloaded May 27, 2022) retrieved from ENSEMBL for gene quantification. The resulting abundance.tsv file served as input for downstream analysis in R 4.1.2. Genes with a total count of <100 reads across all samples were filtered out, leaving a count file with 15,400 reads. This file was normalized and subjected to differential gene expression analysis using DeSeq2 v1.40.2 ^32^. Genes were considered significant if they met the criteria of a p-adjusted value <0.05, log2fold change >2, and a basemean abundance >20 reads. Subsequently, 2,239 genes were filtered from the differentially abundant gene list. These genes were then analyzed for functional pathways using the ShinyGO 0.80 server (http://bioinformatics.sdstate.edu/go/). Four independent biological replicates in each group were analyzed, which yielded consistent results.

### Statistical analysis

The p-values for EAE clinical score were determined using two-way multiple-range analysis of variance (ANOVA) for multiple comparisons. For comparisons involving three groups, one-way ANOVA followed by the Bonferroni post hoc test was utilized. Other p-values were determined using the Mann-Whitney test. Unless specified otherwise, all results are presented as mean ± standard error of the mean (SEM). A p-value of <0.05 was considered statistically significant.

## Results

### Specificity of oligodendroglial lineage mRNA isolation

To purify oligodendroglial lineage-enriched mRNA, we crossed RiboTag mice with the Olig2-cre transgenic mice to get Olig2-cre Rpl22-HA^fl/fl^ mice (refer to as Olig2-cre RiboTag mice). To validate the specific expression of the HA tag in cells of the oligodendroglial lineage, we conducted immunostaining on the spinal cord tissues from Olig2-cre RiboTag mice. We utilized anti-HA antibodies in combination with antibodies targeting GST-π (a mature oligodendrocyte marker), NeuN (a neuronal marker), or GFAP (an astrocyte marker). Our findings revealed that while all GST-π^+^ cells were HA^+^, less than 5% of NeuN^+^ neurons showed HA expression, and no GFAP^+^ astrocytes exhibited HA labeling. This pattern strongly indicates the precise expression of HA within cells of the oligodendroglial lineage (**Fig. 1A**). It had been previously documented ^33^ that olig2-cre expression also occurs in motor neurons, which accounts for the observed HA expression in some neurons within the spinal cord of olig2-cre RiboTag mice. Furthermore, quantitative PCR (Q-PCR) analysis demonstrated an enrichment of oligodendrocyte-specific genes such as MBP, Plp1, and Myrf1 in spinal cord samples following HA mRNA immunoprecipitation from ribosomes of olig2-cre RiboTag mice (**Fig. 1B**). In contrast, there was a notable decrease in the expression of astrocyte-specific Gfap gene and microglial/leukocyte-specific Cd45 gene post immunoprecipitation (**Fig. 1C**), along with a reduction in neuron-specific genes like NeuN, Syn1, and Syt1 (**Fig. 1D**). Collectively, these data support the conclusion that olig2-cre induces HA expression specifically within OLCs, although some expression may also occur in motor neurons.

**Figure 1.**
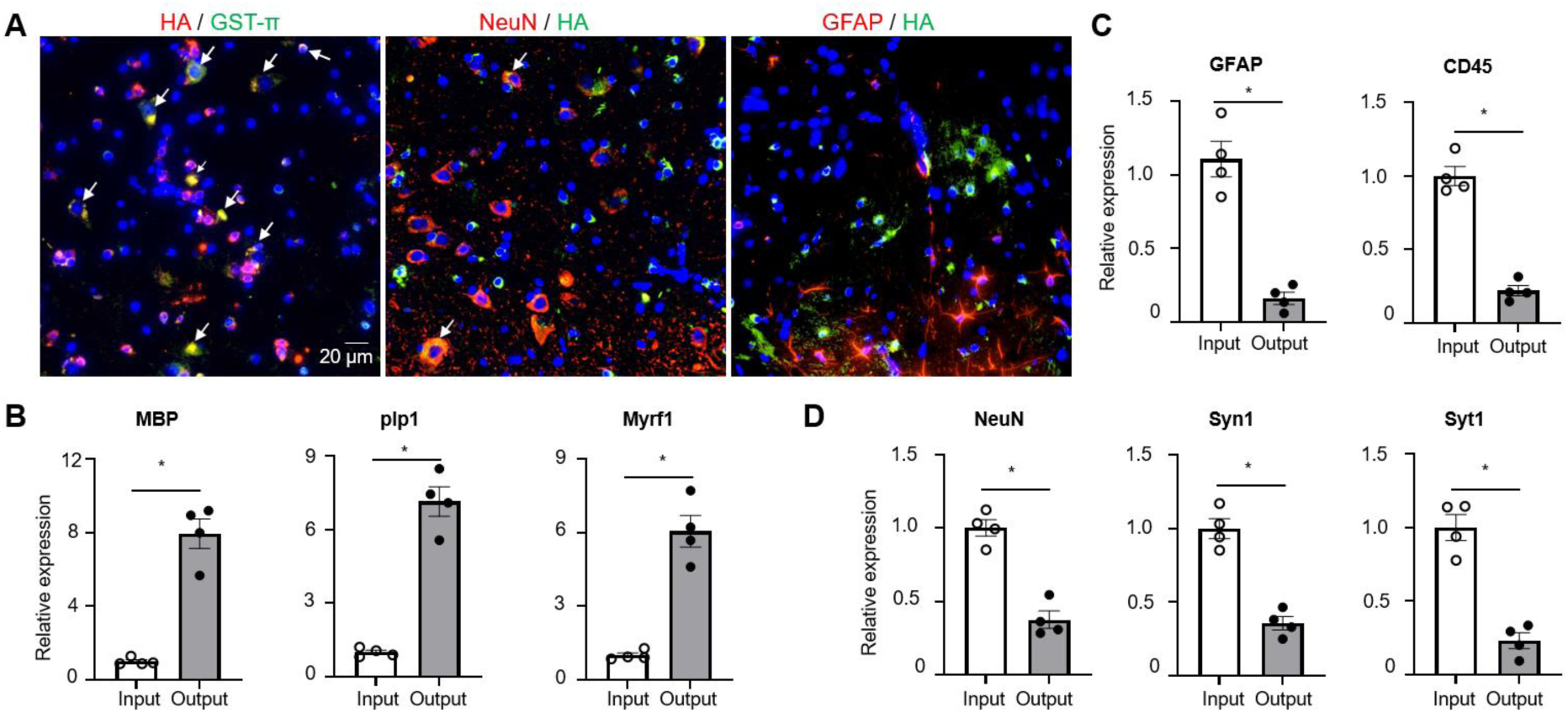
Specificity of oligodendroglial lineage mRNA isolation. (A) Fluorescent immunostaining performed on spinal cords obtained from Oligo2-Cre RiboTag mice using specified antibodies, as indicated. Co-localization is visualized in yellow following image merge, indicated by arrows, with nuclei counterstained using DAPI. (B-D) Q-PCR analysis of transcripts expressed in oligodendrocytes (Panel B), astrocytes, microglia/leukocytes (Panel C), and neuronal cells (Panel D). Total RNA was isolated from spinal cords of naïve Oligo2-Cre RiboTag mice (Input), with output representing mRNA isolated post-immunoprecipitation using anti-HA antibodies from the same group of mice. Gene enrichment or de-enrichment was compared between input and output. n=4/group. Values represent the mean ± SEM, *p<0.05. Data are from one experiment representative of two independent experiments.

### Oligodendroglial lineage reactivity at the onset of EAE

After confirming the specificity of RiboTag-enriched mRNA from OLCs, we proceeded to isolate mRNA from the spinal cords of both naïve Olig2-cre RiboTag mice and Olig2-cre RiboTag mice at the onset of EAE, characterized by tail weakness or loss of tail rigidity. The extracted mRNA underwent bulk RNA-Seq analysis. To interpret the RNA-Seq data, we initiated multi-dimensional scaling (MDS) analysis (**Fig. 2A**) and hierarchical clustering analysis (**Fig. 2B**) using ward D2 distance metrics. Both analyses distinctly segregated data from naïve mice and EAE mice, indicating substantial transcriptomic changes in oligodendroglial cells during EAE pathogenesis. Differential gene expression analysis, employing a Log2 fold change cutoff of 2 and padj=0.05, revealed 1556 upregulated genes and 683 downregulated genes in OLCs from EAE mice compared to naïve controls (**Fig. 2C**). To understand the biological significance of these differentially expressed genes (DEGs), we conducted gene set enrichment analysis (GSEA) utilizing the GO biological process database from the ShinyGO web server. This analysis unveiled the top 20 enriched pathways associated with upregulated genes, including innate and adaptive immune responses, cytokine production, leukocyte activation, and defense responses (**Fig. 2D**). In contrast, the top 20 enriched pathways associated with downregulated genes encompassed neuron development, neuron differentiation, cell morphogenesis, and cytoskeleton organization (**Fig. 2E**), likely implicating neurodegeneration.

**Figure 2.**
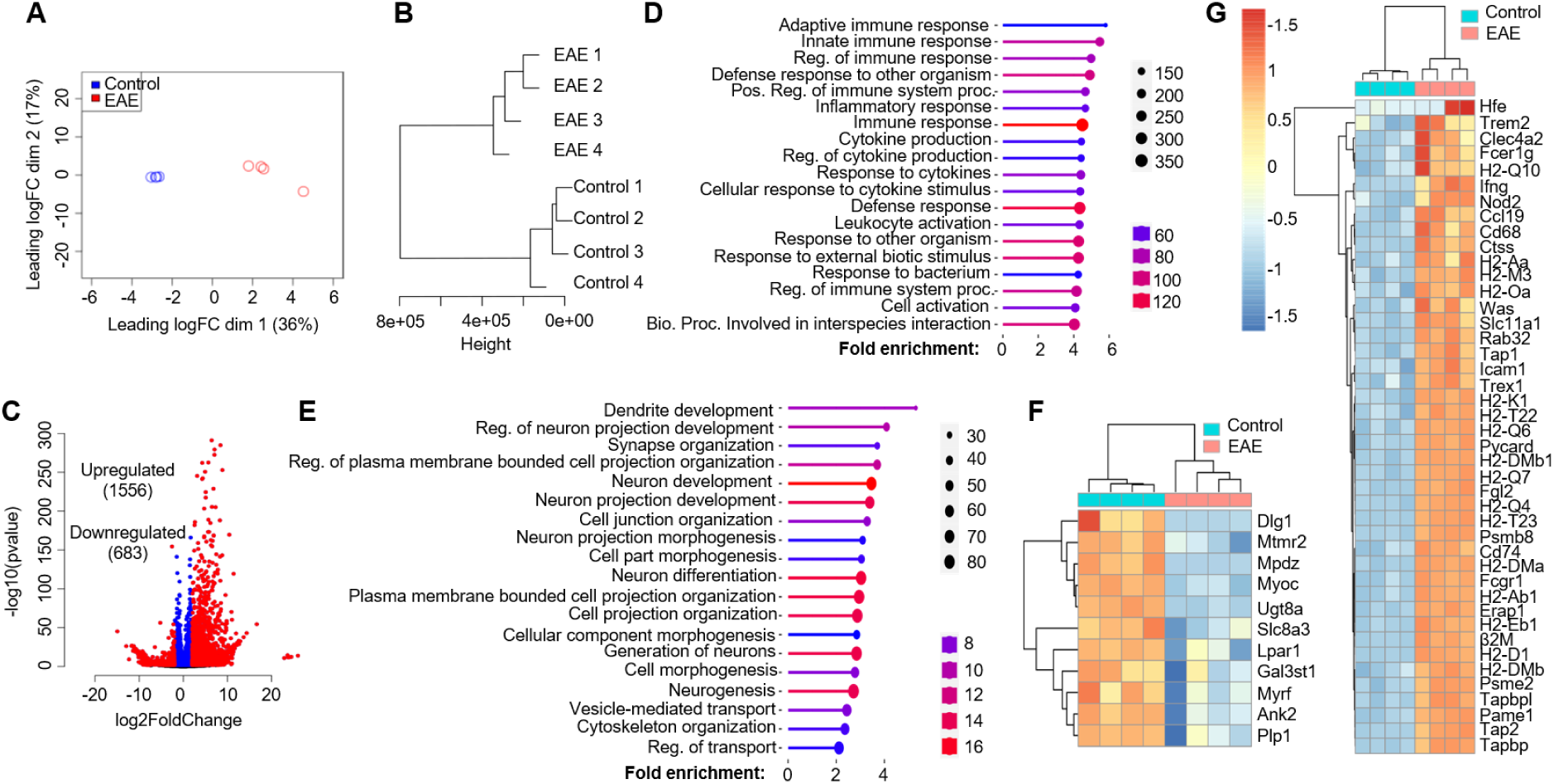
Oligodendroglial lineage reactivity at the onset of EAE. Spinal cords from Oligo2-Cre RiboTag EAE mice at disease onset and naïve control mice were homogenized, and mRNA was isolated following immunoprecipitation with anti-HA antibodies targeting ribosomes. The purified mRNAs underwent RNA-Seq using an Illumina NovaSeq sequencer. The RNA-Seq data were analyzed as follows: (A) Multi-dimensional scaling (MDS) plot displaying RNA-Seq data from spinal cords of Oligo2-Cre RiboTag naïve control mice and EAE mice at disease onset. (B) Hierarchial clustering utilizing ward D2 distance metrics, highlighting distinct clustering of the samples. (C) Identification of differentially expressed genes (DESeq2) in EAE compared to the naïve group, using a Log2Fold change cutoff of 2.0 and padj=0.05. (D) Presentation of the top 20 enriched pathways for 1556 upregulated genes. (E) Top 20 enriched pathways for 683 downregulated genes, as determined by the GO biological process database from the ShinyGO web server. The heatmap illustrates differentially expressed genes categorized into: F) Myelination and Myelin assembly. G) Antigen processing and presentation, within the GO biological process category. n=4/group.

Considering previous single-cell sequencing findings that highlighted the upregulation of genes related to antigen processing and presentation in oligodendrocyte precursor cells (OPCs) during EAE ^18, 19^, we performed gene ontology (GO) analysis. This analysis confirmed a significant upregulation of antigen presenting related genes in EAE mice, such as H2-Q10, H2-Oa, H2-K1, Cd74, H2-Ab1, H2-Eb1, Tap1, and Tap2, among others (**Fig. 2G**). Additionally, GO analysis revealed a decrease in the expression of myelination-related genes, including Ugt8a, Myrf, and Plp1, suggesting demyelination in EAE mice (**Fig. 2F**).Furthermore, in line with GSEA analysis indicating enriched cytokine production, response to cytokines, and innate immune response, we conducted GO analysis specifically focusing on cytokine receptors and toll-like receptors (**Fig. S1A**), chemokines and chemokine receptors (**Fig. S1B**), IFN-β response genes (**Fig. S1C**), and IFN-γ response genes (**Fig. S1D**). Interestingly, we observed a substantial upregulation of type I and type-II interferon response genes in EAE OLCs (**Fig. S1E**). Notably, our GO analysis confirmed the expression of known cytokine receptors like Il1r1, Ifnar1, Il4ra, and Il17ra in OLCs, validating our RiboTag RNA-Seq dataset (**Fig.S1A**). Moreover, we identified new receptor expressions in oligodendroglial cells, such as Csf2r, Il3ra, Il13ra1, and Il18r1, among others (**Fig.S1A-B**). Overall, our RNA-Seq dataset indicates a comprehensive inflammatory signature in OLCs, suggesting their potential proinflammatory role in autoimmune encephalomyelitis.

### Validation of RiboTag RNA-Seq data via qRT-PCR

The RNA-Seq data provided intriguing insights; however, it was imperative to validate the gene expressions in Oligodendrocyte Lineage Cells (OLCs) in the context of Experimental Autoimmune Encephalomyelitis (EAE). To achieve this, we utilized purified mRNA from OLCs of both Olig2-cre RiboTag naïve mice and Olig2-cre RiboTag EAE mice, employing SYBR Green Real-time PCR Master Mix for qRT-PCR analysis. Initially, we confirmed that genes associated with myelination, such as Plp1, Mbp, Ugt8a, Myrf, and Olig2, exhibited a significant decrease in EAE OLCs compared to their expression levels in naïve OLCs. Concurrently, the neuronal marker gene Syn1 also displayed reduced expression, indicating demyelination and degeneration in EAE mice (**Fig. 3A**). Furthermore, our validation revealed the upregulation of key antigen processing and presenting genes in EAE OLCs, including Citta, CD74, Nlrc5, H2-K1, Tbp1, and H2-Aa (**Fig. 3B**), thereby supporting the hypothesis that OLCs, particularly Oligodendrocyte Progenitor Cells (OPCs), may function as antigen-presenting cells in MS/EAE ^11, 19, 34^. Lastly, we confirmed the increased expression of Ifngr1 and Ifnar2 in EAE OLCs, along with elevated levels of interferon response genes such as ISG56, Ifit2, Ccl2, and Cxcl10 (**Fig. 3C**). Collectively, our Q-PCR analysis validated numerous gene expressions in EAE OLCs as observed in the RNA-Seq dataset, thus affirming the reliability of the RNA-Seq data.

**Figure 3.**
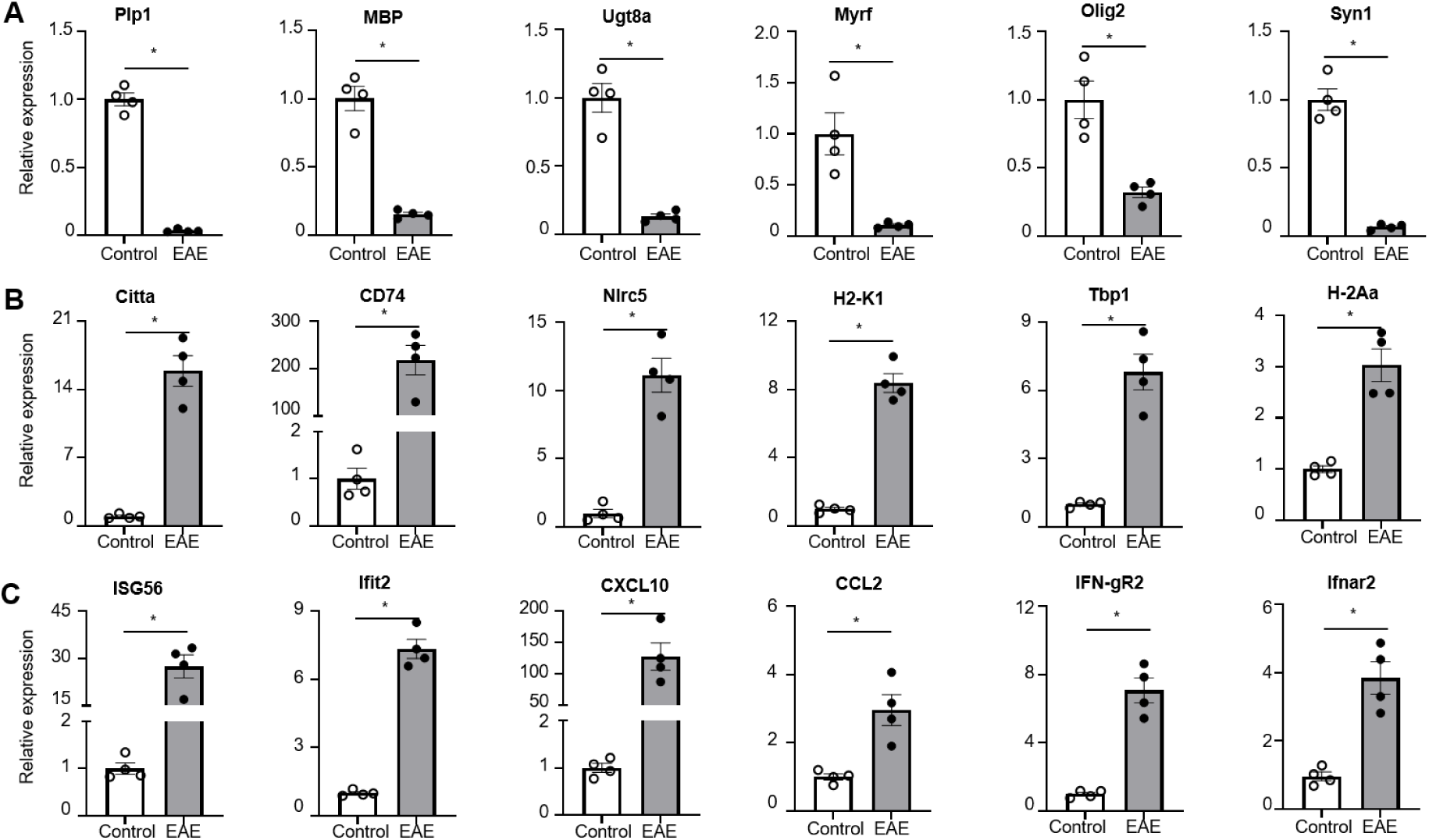
Q-PCR validation of RiboTag RNA-Seq data. Spinal cords from Oligo2-Cre RiboTag EAE mice at disease onset and naïve control mice were homogenized, and mRNA was isolated following immunoprecipitation with anti-HA antibodies targeting ribosomes. The purified mRNAs underwent Q-PCR analysis to validate RNA-Seq data from Figure 2. (A) Demonstrates decreased expression of myelination-related genes in the EAE group, as indicated. (B) Shows upregulation of genes associated with antigen processing and presentation in the EAE group. (C) Indicates upregulation of interferon response genes in the EAE group. Values represent the mean ± SEM, *p<0.05. Data are from one experiment representative of two independent experiments.

### Cell-intrinsic IFN-γ signaling in oligodendroglial lineage promotes EAE pathogenesis

The RNA-Seq data from OLCs revealed robust activation of interferon response genes, notably IFN-γ and IFN-β response genes. This indicates a potential cell-intrinsic role of IFN-γ and IFN-β signaling within the oligodendroglial lineage, which may significantly contribute to the pathogenesis of experimental autoimmune encephalomyelitis (EAE). Furthermore, the upregulation of IFNGR and IFNAR in OLCs at the onset of EAE was validated through qRT-PCR, confirming the initial observations (**Fig.3**). The role of IFN-γ in the context of multiple sclerosis (MS) and EAE is intricate. On one hand, administering IFN-γ to MS patients and EAE mice has been shown to exacerbate central nervous system (CNS) inflammation and clinical symptoms ^35–37^. Conversely, studies on IFN-γ and IFNGR knockout mice indicate increased susceptibility to EAE, resulting in heightened morbidity and mortality ^38–40^. Adding to the complexity, IFN-γ exhibits a stage-specific dual role in EAE. Treatment with IFN-γ during the initiation phase of EAE exacerbates disease severity, whereas administration during the effector phase mitigates the disease ^41^. Moreover, modulation of IFN-γ signaling yields distinct effects in different CNS cell types. Silencing IFN-γ signaling in astrocytes alleviates EAE, whereas its suppression in microglia exacerbates the disease ^42, 43^. Despite these findings, the specific impact of cell-intrinsic IFN-γ signaling derived from OLCs on EAE pathogenesis remains unclear. We hypothesized that IFN-γ signaling within OLCs contributes to the exacerbation of EAE pathology.

To test this hypothesis, we conducted crossbreeding involving Ifngr1^fl/fl^ mice and Olig2-Cre Ifngr1^fl/fl^ mice, resulting in the generation of Olig2-Cre Ifngr1^fl/fl^ conditional knockout mice (referred to as IFNGR cKO) and Ifngr1^fl/fl^ wild-type control mice (referred to as IFNGR WT). Initially, we confirmed PCR genotyping accuracy through flow cytometry analysis of IFNGR in OPCs. We cultured OPCs from embryonic pups of both Ifngr1^fl/fl^ and Olig2-Cre Ifngr1^fl/fl^ genotypes and stained them with OPC marker CD140a antibody and IFNGR antibody. The flow cytometry data indicated successful ablation of IFNGR in OPCs from IFNGR cKO mice (**Fig. 4A, right**), whereas IFNGR expression was confirmed in IFNGR WT mice (**Fig. 4A, left**), consistent with our RNA-Seq and Q-PCR findings. Subsequently, both IFNGR cKO mice and IFNGR WT control mice were subjected to active EAE induction via subcutaneous immunization with MOG^35–55^ peptides. Notably, IFNGR cKO mice exhibited significantly lower EAE clinical scores compared to WT control mice (**Fig. 4B**), indicative of attenuated disease severity. This was further supported by markedly lower maximum scores and cumulative scores in IFNGR cKO mice, highlighting the reduced severity of EAE (**Fig. 4C**). Interestingly, we did not observe significant differences in disease onset, time to peak, or disease incidence between the two groups (**Fig. 4C**), suggesting that intrinsic IFN-γ signaling in OLCs primarily impacts the effector stage of EAE. Consistent with the amelioration of disease severity, histopathological analysis revealed reduced accumulation of infiltrating immune cells and resultant demyelination in the spinal cord of IFNGR cKO mice compared to controls (**Fig. 4D**). Collectively, these findings strongly suggest that OLCs-intrinsic IFN-γ signaling plays a crucial role in promoting EAE pathogenesis.

**Figure 4.**
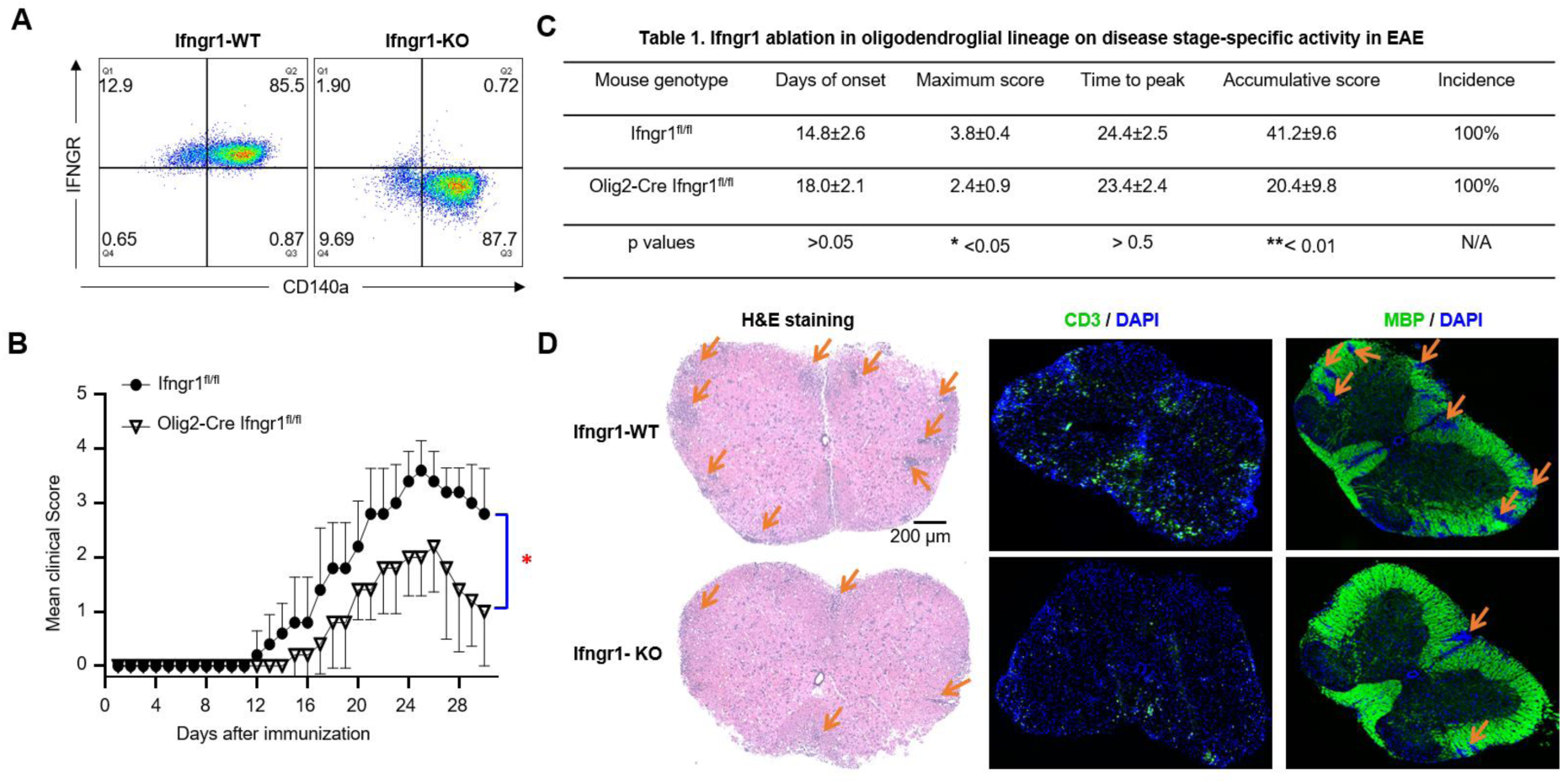
Cell-intrinsic IFN-γ induced signaling in oligodendroglial lineage promotes EAE pathogenesis. (A) Flow cytometry analysis indicates Ifngr1 deletion in OPCs cultured from Olig2-Cre Ifngr1^fl/fl^ mice as indicated. OPCs were cultured from embryonic pups of Ifngr1^fl/fl^ mice and Olig2-Cre Ifngr1^fl/fl^ genotypes. (B) Mean clinical score of EAE in Ifngr1^fl/fl^ mice and Olig2-Cre Ifngr1^fl/fl^ mice induced by active immunization with MOG_35–55_ (p < 0.05, two-way ANOVA). (C) Disease stage-specific activities in EAE influenced by specific Ifngr1 ablation in oligodendroglial lineage cells. (D) Hematoxylin and eosin (H&E) staining, anti-CD3 immunostaining, and myelin basic protein (MBP) staining were performed as indicated on lumbar spinal cords harvested 25 days after immunization. Data are from one experiment representative of three independent experiments, with n = 5/group in each experiment. Error bars indicate SEM; *p < 0.05, **p < 0.01.

### Cell-intrinsic IFN-β signaling in oligodendroglial lineage is dispensable for EAE pathogenesis

In contrast to the complex role of IFN-γ in the development of MS/EAE, systemic administration of IFN-β has emerged as the most widely used therapy for MS, offering prolonged periods of remission, reduced severity of relapses, and decreased inflammatory lesions in the CNS ^44–47^. The absence of either IFN-β or IFNAR has been linked to a more severe and chronic form of EAE ^48–50^. However, the underlying mechanisms behind the therapeutic effects of IFN-β on MS/EAE remain unclear. This prompted us to investigate whether intrinsic IFN-β signaling in OLCs acts as a negative regulator in EAE pathogenesis.

To address this question, we initially attempted to obtain Ifnar1^fl/fl^ and Olig2-Cre Ifnar1^fl/fl^ mice for EAE experiments. However, we encountered difficulties in obtaining Olig2-Cre Ifnar1^fl/fl^ mice due to unknown reasons (data not shown). As an alternative, we utilized Pdgfra-cre^ER^ Ifnar1^fl/fl^ mice. We verified the deletion efficiency by culturing OPCs from Pdgfra-cre^ER^ Ifnar1^fl/fl^ mice and inducing Ifnar1 deletion through 4-OHT treatment (an active metabolite of tamoxifen) in these cells. Our results showed more than 95% deletion of Ifnar1 in 4-OHT-treated OPCs compared to vehicle-treated OPCs (**Fig. 5A**), affirming the effectiveness of cre in mediating IFNAR ablation in OPCs.

**Figure 5.**
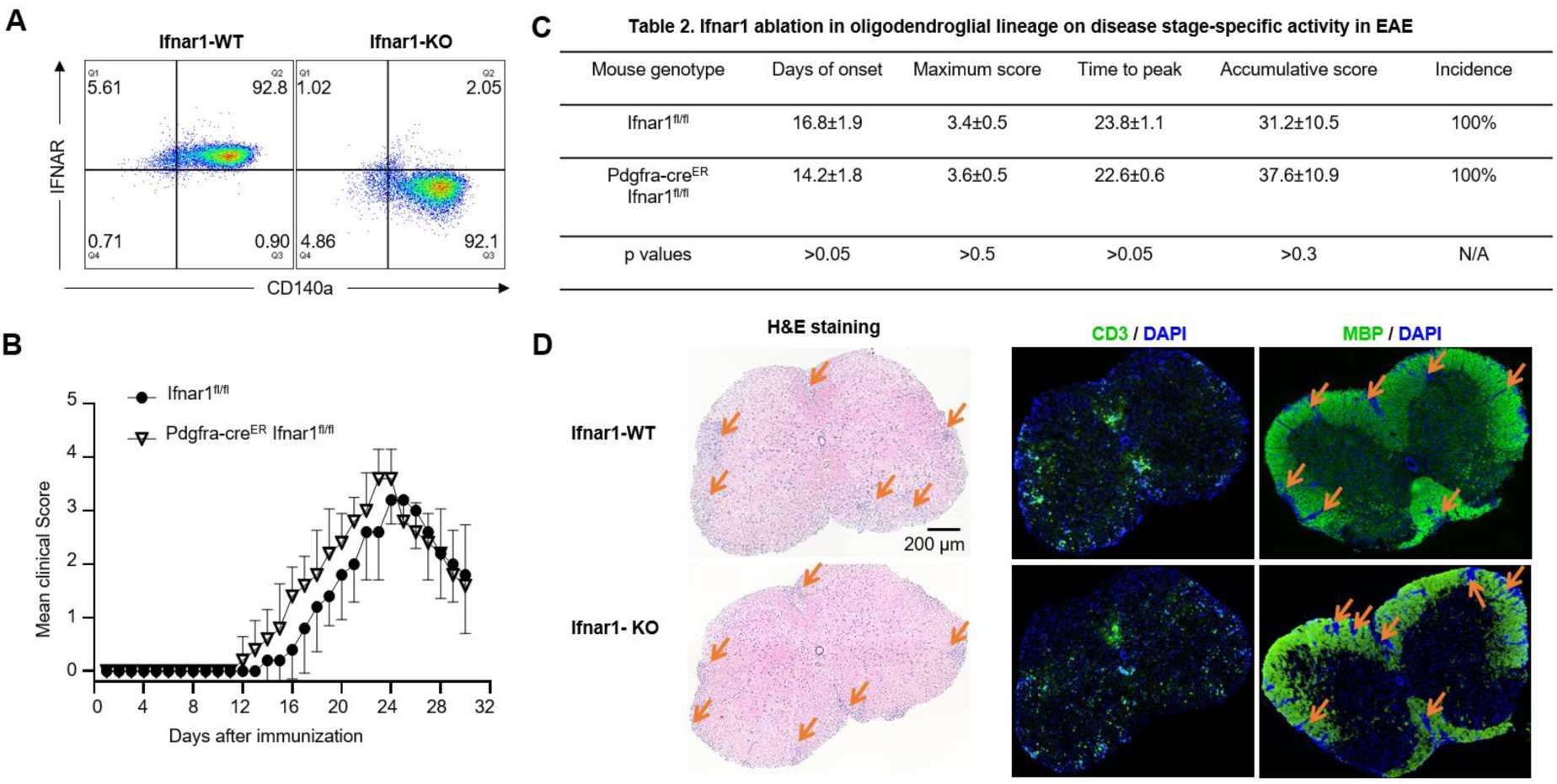
Cell-intrinsic IFN-β induced signaling in oligodendroglial lineage is dispensable EAE pathogenesis. (A) Flow cytometry analysis indicates Ifnar1 deletion in OPCs cultured from Pdgfra-cre^ER^ Ifnar1^fl/fl^ mice after 4-Hydroxytamoxifen (4-OHT) induction as indicated. OPCs were cultured from embryonic pups of Ifngr1^fl/fl^ mice and Pdgfra-Cre^ER^ Ifnar1^fl/fl^ genotypes, 1μM OHT was used to induce Cre expression and Ifnar1 deletion. (B) Mean clinical score of EAE in Ifnar1^fl/fl^ mice and Pdgfra-cre^ER^ Ifnar1^fl/fl^ mice induced by active immunization with MOG_35–55_ (p < 0.05, two-way ANOVA). (C) Disease stage-specific activities in EAE influenced by specific Ifnar1 ablation in oligodendroglial lineage cells. (D) Hematoxylin and eosin (H&E) staining, anti-CD3 immunostaining, and myelin basic protein (MBP) staining were performed as indicated on lumbar spinal cords harvested 25 days after immunization. Data are from one experiment representative of three independent experiments, with n = 5/group in each experiment. Error bars indicate SEM; *p < 0.05.

Subsequently, we induced Ifnar1 deletion in OLCs of Pdgfra-cre^ER^ Ifnar1^fl/fl^ mice in vivo using tamoxifen. The control group consisted of Ifnar1^fl/fl^ littermate mice treated with tamoxifen. Henceforth, the tamoxifen-treated Pdgfra-cre^ER^ Ifnar1^fl/fl^ mice are denoted as IFN-AR cKO mice, while the tamoxifen-treated Ifnar1^fl/fl^ mice are labeled as IFNAR WT control mice. Both groups underwent active EAE induction. Surprisingly, we did not observe a significant difference in mean EAE clinical scores between the two groups post-EAE induction (**Fig. 5B**). Consistently, there were no discernible differences in disease onset, maximum clinical score, time to peak, cumulative clinical score, or disease incidence between the experimental groups (**Fig. 5C**). Corresponding to the disease severity observed between the groups, histopathological analysis revealed comparable accumulation of infiltrating immune cells and resultant demyelination in the spinal cord of IFN-AR cKO mice compared to controls (**Fig. 5D**). Consequently, we conclude that cell-intrinsic IFN-β signaling in OLCs is dispensable for the pathogenesis of EAE.

### Differential immune responses of OPCs to IFN-γ and IFN-β

The distinct impacts of IFN-γ and IFN-β signaling within OLCs are pivotal in understanding EAE pathogenesis, prompting an exploration into their disparate effects. Oligodendrocytes treated with IFN-γ experience apoptosis or necrosis ^51, 52^. IFN-γ can initiate death related pathway in OPCs depending on the dose ^53^, but it can also suppress OPC differentiation and maturation without overt death at optimal dosage ^54^. Notably, IFN-γ inhibits cell cycle exit during OPC differentiation, rendering them susceptible to apoptosis and hindering their maturation ^55^. Conversely, IFN-β exerts no discernible influence on OPC proliferation, differentiation, or survival, indicating a differential impact compared to IFN-γ ^56^. Given that demyelination characterizes MS/EAE, these findings partially elucidate why intrinsic IFN-γ signaling in OLCs, as opposed to IFN-β, contributes to EAE pathogenesis, albeit without considering the inflammatory context of oligodendroglia in these earlier studies.

To analyze the targeted gene expression by IFN-γ and IFN-β at the onset of EAE, we conducted GO analysis specifically focusing on IFN-γ and IFN-β response genes from our RiboTag RNA-seq dataset. We found a specific enrichment of IFN-γ and IFN-β response immune genes in OLCs at EAE onset, revealing both shared and distinct gene expression patterns (**Fig.S1C-E**). To delve deeper, we conducted qRT-PCR assays in vitro to decipher how OPCs respond differentially to IFN-γ and IFN-β treatments. Our findings indicate a robust response of genes related to MHC class I antigen processing and presentation (Tap1, β2m, and H2-D1) to IFN-γ compared to IFN-β (**Fig.6A**). Similarly, genes linked to MHC class II antigen processing and presentation (H2-Aa, H2-Eb1, and Cd74) showed significantly higher upregulation in response to IFN-γ (**Fig.6B**). Further analysis revealed a unique set of genes (Nlrc5, Citta, Ccl2, and Ccl7) that responded specifically to IFN-γ, consistent with our RNA-Seq data (**Fig. 6C**). Intriguingly, IFN-β, but not IFN-γ, induced the expression of Irf1 and Irf7critical for host defense mechanisms in OPCs (**Fig.6D**). These findings collectively suggest that OLCs-intrinsic IFN-γ signaling may exacerbate EAE pathogenesis by enhancing antigen processing, presentation, and chemokine production (e.g., Ccl2 and Ccl7), while OLCs-intrinsic IFN-β signaling likely contributes to host defense mechanisms.

**Figure 6.**
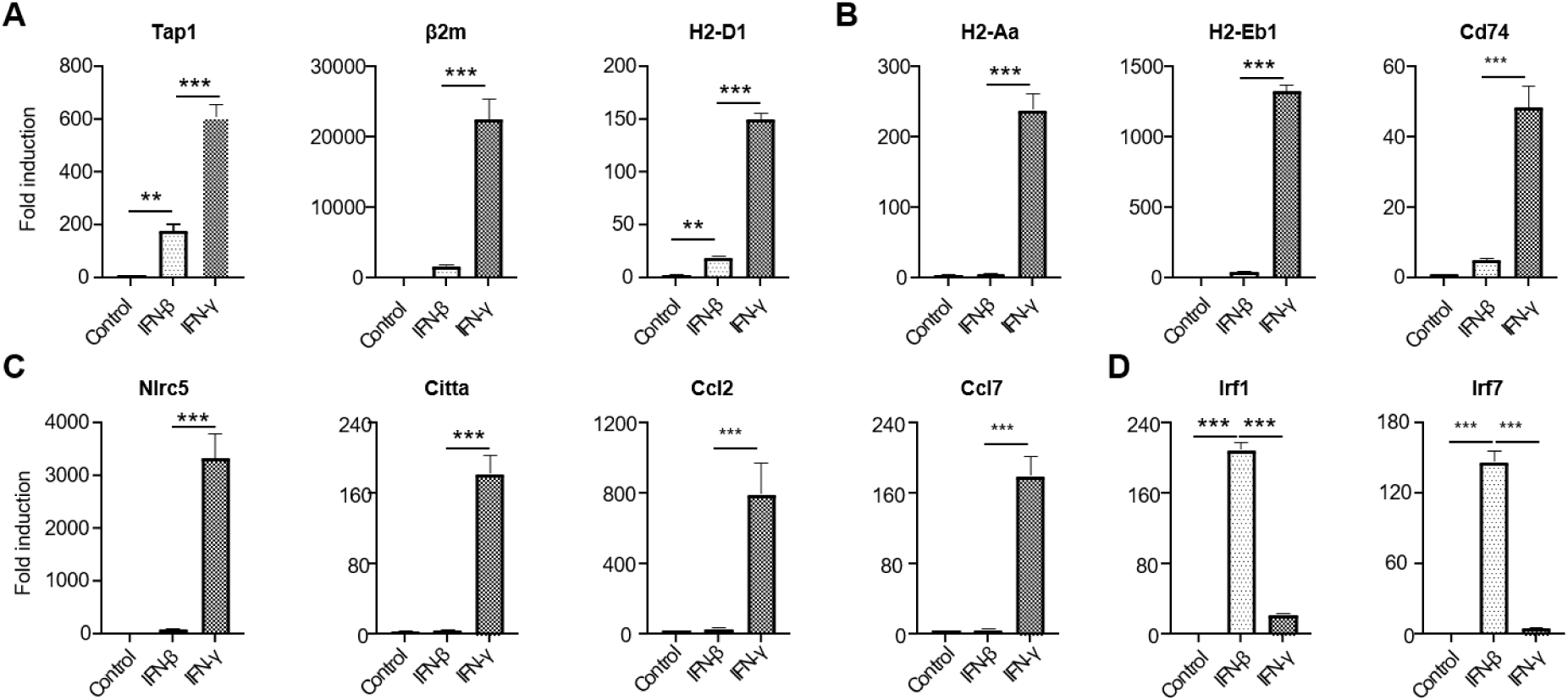
Differential roles of type I and type II interferons on OPC. Oligodendrocyte progenitor cells (OPCs) were cultured from embryonic pups of C57BL/6 mice, following the procedures outlined in the Material and Methods section. The OPCs were then divided into three groups: untreated, treated with 20 ng/ml IFN-β, or treated with 20 ng/ml IFN-γ, respectively, for a duration of 6 hours. Subsequently, mRNA was extracted for Q-PCR analysis. (A) Depicts the expression of MHC-I-related genes. (B) Depicts the expression of MHC-II-related genes. (C) Highlights genes specifically induced by IFN-γ. (D) Highlights genes specifically induced by IFN-β. Data are from one experiment representative of three independent experiments, with n=3 technical repeats per group in each experiment. Error bars represent SEM, with ***p<0.001.

## Discussion

The RiboTag approach, which targets ribosome-associated mRNAs, provides a more accurate representation of actively translated genes, offering a clearer view of the translational state within specific cellular compartments. Thus, mRNAs isolated through the RiboTag method reflect the translatome, providing a snapshot of gene expression during active translation ^24, 57^. Using RiboTag RNA-Seq on EAE mice at disease onset, we identified 1,556 upregulated and 683 downregulated genes in OLCs compared to healthy controls. The upregulated genes were primarily involved in innate and adaptive immune responses, particularly cytokine production and defense responses. In contrast, downregulated genes were mainly linked to neuron development, cell morphogenesis, regulation of transport, and myelination. Among the enriched pathways, IFN-γ and IFN-β responsive genes were highly represented, prompting us to investigate their OLC-intrinsic roles in EAE pathogenesis. Notably, we found a significant pathogenic role for cell-intrinsic IFN-γ signaling in OLCs, whereas IFN-β signaling appeared dispensable for disease progression or suppression.

Our study reveals a pro-inflammatory role for OLCs in EAE, suggesting their potential involvement in MS. Using RiboTag RNA-Seq, we observed a distinct transcriptional shift in OLCs during EAE, which aligns with immune modulation. Our data suggest that OLCs play an active role in neuroinflammation, in addition to their well-established function in myelination. This finding supports previous work by the Castelo-Branco group, who demonstrated immune gene expression in both OPCs and mature oligodendrocytes in EAE mice using single-cell RNA sequencing (scRNA-Seq), referring to these as inflammatory oligodendrocyte lineage cells (iOLCs) ^18^. They identified upregulated genes associated with antigen processing, T cell cytotoxicity, immunoglobulin secretion, cytokine production (e.g., IL-4, IL-13, IL-8), dendritic cell differentiation, and interferon response genes in both OPCs and mature oligodendrocytes ^18^. Our analysis supports these findings and expands upon them, identifying additional cytokines, chemokines, and their receptors, as well as various toll-like receptors in OLCs. Further comparison of our RiboTag data with single-cell data could offer insights into the methodological advantages and the specific contributions of OPCs and mature oligodendrocytes to OLC-mediated neuroinflammation in EAE.

In this study, we primarily validated the sequencing data in OPCs, as they are easier to culture in vitro and can be identified by surface markers like NG2 and CD140a/PDGFRα. Future work will extend validation to mature oligodendrocytes using reporter mice. Altogether, our findings provide compelling evidence that oligodendroglial lineage cells play a significant role in EAE pathogenesis, an animal model for MS. By leveraging RiboTag technology, we were able to isolate and analyze mRNA specifically from these cells, revealing a complex and dynamic transcriptomic landscape during disease progression.

Although Olig2 is a well-established lineage marker for oligodendrocytes, previous studies have shown that Olig2-Cre is also partially expressed in motor neurons ^33^. Our histology analysis similarly showed sparse HA tag expression in neurons. Interestingly, downregulated genes in our dataset were largely associated with neuron development, cell morphogenesis, transport regulation, and myelination. This raises the possibility that some downregulated genes linked to neuronal development or function may originate from neurons rather than OLCs. However, the situation appears more complex, as these pathways are also implicated in scRNA-Seq datasets from OLCs ^18^, suggesting that OLCs share some gene expression patterns with neurons in EAE, possibly even under normal conditions. The downregulation of genes involved in neuron development and myelination supports the idea that CNS inflammation disrupts normal oligodendrocyte function, contributing to the demyelination observed in MS. Validation of these findings through qRT-PCR strengthens the reliability of the RNA-Seq data and underscores the importance of OLCs within the inflammatory environment of EAE. Future studies should further investigate these shared gene expression patterns to deepen our understanding of OLC and neuron interactions in disease and homeostasis.

Both RiboTag RNA-Seq and scRNA-Seq data indicate an enriched response to IFN-γ and IFN-β in OLCs during EAE ^18^. In vitro studies have demonstrated that IFN-γ promotes antigen processing and presentation in OPCs and activates co-cultured T cells, including both CD4+ and CD8+ subsets ^18, 19^. Additionally, IFN-γ inhibits OPC maturation and proliferation in vitro ^55^. However, it remains unclear whether IFN-γ exerts any cell-intrinsic effects in OLCs that influence the pathogenesis of EAE and MS. To address this, we specifically deleted IFNGR1 in OLCs using Olig2 Cre mice and assessed the impact on EAE progression. Our data indicates that IFN-γ signaling within OLCs contributes to the pathogenesis of EAE. The role of IFN-γ in EAE/MS remains controversial, with studies suggesting both beneficial and pathogenic effects. While IFN-γ administration in MS patients and EAE mice exacerbates neuroinflammation and clinical symptoms ^35–37^, IFN-γ and IFNGR knockout mice exhibit increased susceptibility to EAE, with heightened morbidity and mortality ^38–40^. Furthermore, IFN-γ has stage-specific effects on EAE development ^41^. Compounding the complexity, modulation of IFN-γ signaling has distinct outcomes depending on the CNS cell type. For instance, silencing IFN-γ signaling in astrocytes alleviates EAE, while its suppression in microglia exacerbates the disease ^42^. Our findings extend this concept to OLCs, highlighting the importance of targeted therapies in specific cellular compartments during EAE/MS. Additionally, Th17 cells are critical to the pathogenesis of EAE/MS. Our group previously showed that IL-17 produced by Th17 cells drives EAE progression by targeting OPCs. Interestingly, fate mapping of Th17 cells in EAE revealed that autoantigen-specific Th17 cells convert to IL-17^+^IFN-γ^+^ “Ex-Th17 cells” as the disease progresses ^58^. Thus, our findings provide new insight into why Th17 cells are pivotal in EAE pathogenesis: they produce both IL-17 and IFN-γ, and target OLCs in the process.

While previous studies suggest that IFN-β does not significantly influence OPC proliferation, differentiation, or survival, indicating a differential effect compared to IFN-γ ^56^, our preliminary findings suggest that OPCs do respond to IFN-β by expressing IFN-stimulated genes. IFN-β is a first-line treatment for relapsing forms of multiple sclerosis (MS), and the absence of either IFN-β or its receptor (IFNAR) has been linked to more severe and chronic forms of EAE ^48–50^. Furthermore, the engagement of IFNAR on myeloid cells exacerbates EAE, leading to an enhanced effector phase and increased mortality ^59^. However, the cell-intrinsic role of IFN-β in OLCs during EAE remains unclear. Our data suggests that this intrinsic pathway may be dispensable for EAE development. Interestingly, loss of neuronal IFN-β-IFNAR signaling has been associated with Parkinson’s disease-like dementia ^60^, suggesting a protective role for IFN-β in neuronal homeostasis. In our study, we were unable to obtain Olig2Cre Ifnar1^fl/fl^ pups, although Olig2Cre Ifnar1^fl/+^ mice appeared normal, suggesting that IFN-β-IFNAR signaling may play a role in OLC development and warrants further investigation. In practice, distinguishing between type I and type II interferon response genes remains a challenge ^61^. To better understand the differential roles of OLC-intrinsic IFN-γ and IFN-β signaling in EAE pathogenesis, we identified genes related to antigen processing and presentation that showed elevated expression in response to IFN-γ. Additionally, a unique set of genes (Nlrc5, Citta, Ccl2, and Ccl7) was specifically induced by IFN-γ in OPCs, consistent with our RNA-Seq data. In contrast, IFN-β induced a much higher expression of Irf1 and Irf7, both of which are critical for host defense mechanisms. Further elucidation of the distinct responses to IFN-γ and IFN-β in OLCs in vivo will enhance our understanding of interferon-stimulated genes (ISGs) and their potential as therapeutic targets in MS.

In conclusion, our data highlight oligodendroglial lineage cells as active contributors to the immune response in MS. Targeting cell-specific inflammatory pathways within OLCs could represent a novel approach for managing neuroinflammation and promoting CNS repair in MS. Further studies exploring the modulation of OLC-specific cytokine signaling and its effects on disease progression could pave the way for innovative therapeutic strategies.

## Authors’ Contributions

Y.W. performed most of the experiments, interpreted the data, wrote and reviewed the manuscript. S.G. analyzed sequencing data and contributed to manuscript writing. A.M. contributed to data interpretation and manuscript revision. Z.K. was integral for experimental design, manuscript writing, data interpretation and project coordination.

---

**Supplemental Figure 1.**
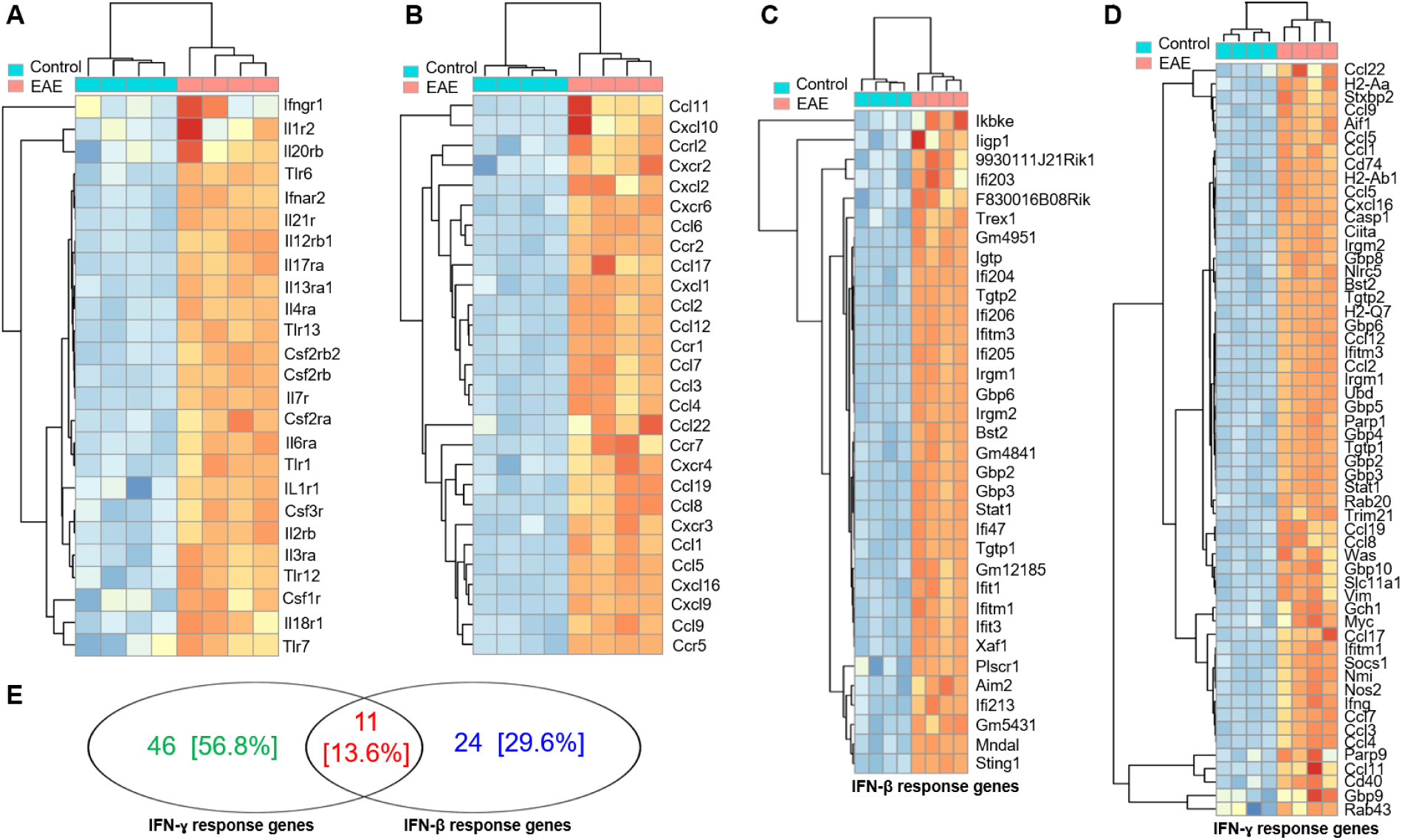
Heatmap illustration of differentially expressed genes. Spinal cords from Oligo2-Cre RiboTag EAE mice at disease onset and naïve control mice were homogenized, and mRNA was isolated via immunoprecipitation with anti-HA antibodies targeting ribosomes. The purified mRNAs underwent RNA-Seq using an Illumina NovaSeq sequencer, and the resulting data were analyzed. The heatmap depicts differentially expressed genes categorized as follows: A) Cytokine receptors and Toll-like receptors (Tlrs). B) Chemokine and chemokine receptors. C) IFN-β response genes. D) IFN-γ response genes. Within the GO biological process category (Panel A-D). E) Venn diagram of IFN-γ and IFN-β response genes. n = 4/group. Related to Figure 2.

